# Melting dsDNA donor molecules potentiates precision genome editing in *C. elegans*

**DOI:** 10.1101/2020.08.03.235036

**Authors:** Krishna S. Ghanta, Craig C. Mello

## Abstract

CRISPR genome editing has revolutionized genetics in many organisms. In the nematode *Caenorhabditis elegans* one injection into each of the two gonad arms of an adult hermaphrodite exposes hundreds of meiotic germ cells to editing mixtures, permitting the recovery of multiple indels or small precision edits from each successfully injected animal. Unfortunately, particularly for long insertions, editing efficiencies can vary widely, necessitating multiple injections, and often requiring co-selection strategies. Here we show that melting double stranded DNA (dsDNA) donor molecules prior to injection increases the frequency of precise homology-directed repair (HDR) by several fold for longer edits. We describe troubleshooting strategies that enable consistently high editing efficiencies resulting, for example, in up to 100 independent GFP knock-ins from a single injected animal. These efficiencies make *C. elegans* by far the easiest metazoan to genome edit, removing barriers to the use and adoption of this facile system as a model for understanding animal biology.

---

In the nematode worm *C. elegans*, genome editing can be achieved by direct injection of Cas9 guide-RNA ribonucleoprotein (RNP) complexes into the syncytial ovary (Cho *et al*. 2013; Paix *et al*. 2015; Dokshin *et al*. 2018). In the worm germline, such injections afford the editing machinery simultaneous access to hundreds of meiotic germ nuclei that share a common cytoplasm. Under optimal conditions the frequency of F1 progeny with indels caused by non-homologous end joining (NHEJ) can be greater than 90% of those progeny expressing a co-injection plasmid marker gene (Dokshin *et al*. 2018). Leveraging these high cutting efficiencies, precise genome editing is readily achieved using short (under ∼200 nucleotide [nt]), single-stranded oligodeoxynucleotide (ssODN) donors, permitting insertions of up to ∼150 nt in length (Arribere *et al*. 2014; Zhao *et al*. 2014; Paix *et al*. 2015; Prior *et al*. 2017; Dokshin *et al*. 2018). However, with longer dsDNA donors (∼1kb), HDR events are recovered at lower frequencies, require more complex protocols, high concentrations of the donor DNA, and typically require screening the broods of multiple injected animals (Tzur *et al*. 2013; Arribere *et al*. 2014; Kim *et al*. 2014; Ward 2015; Paix *et al*. 2016; Schwartz and Jorgensen 2016; Paix *et al*. 2017; Dokshin *et al*. 2018; Farboud *et al*. 2019; Silva-Garcia *et al*. 2019; Vicencio *et al*. 2019).

There are multiple reasons why longer repair templates may be less efficient as donors for HDR compared to ssODNs. First, empirical studies suggest that long dsDNA is more toxic than short single-stranded DNA (Mello *et al*. 1991), limiting safe donor concentrations to less than 200 ng/µl for ∼1kb donors. Second, upon injection into germline cytoplasm, dsDNA molecules quickly form large extra-chromosomal arrays via both end-joining and homologous recombination pathways, and appear to do so while sequestered away from genomic DNA (Stinchcomb *et al*. 1985; Mello *et al*. 1991). Concatenation of donor molecules into large arrays would have the effect of lowering the number of individual molecules available to access and to template repair at the target site double strand break (DSB). Moreover, if injected DNA assembles concatenates while sequestered from the nuclear DNA—perhaps within de novo nucleus-like organelles (Forbes *et al*. 1983)—this process could preclude templated repair of a genomic target site until after the sequestered concatenates gain nuclear access after nuclear envelope breakdown occurs post-fertilization.

In a recent study, we showed that CRISPR-mediated HDR could be increased ∼4-fold by mixing, melting, and re-annealing overlapping donor molecules to create donor templates with single-stranded overhangs (Dokshin *et al*. 2018). In those previous studies, we limited our analysis to a cohort of F1 ‘Roller’ progeny that express the co-injection marker gene *rol-6 (su1006)*. Here, to explore editing efficiency outside the Roller cohort, we scored the entire brood of each injected animal for precisely edited progeny that incorporate and express fluorescent protein markers (GFP or mCherry). We show that the vast majority of insertions occurred later in the brood, after the cohort of progeny that express the Roller phenotype. Whereas overhangs improved the frequency of editing among the F1 Rollers (Dokshin *et al*. 2018), they had no benefit within this latter segment of the brood. Instead, melting the donor molecules, alone, sufficed to increase the HDR frequency to as high as 50% of the post-injection progeny. We provide a protocol and troubleshooting strategies that enable even a novice injector to achieve their editing goals and to optimize editing efficiencies.

## MATERIALS AND METHODS

Detailed editing protocol is provided in Supplemental Material, **File S1**.

### Strains and genetics

All the strains were generated in the Bristol N2 background unless specified otherwise and cultured on normal growth media (NGM) plates seeded with OP50 bacteria (Brenner 1974). Strains used in this study are listed in Supplemental Material, **Table S1**.

At CSR-1 locus, GFP was introduced between FLAG::linker(9bp) and TEV in FLAG::linker::TEV::CSR-1 strain.

### Scoring methodology

Injected P0 animals were individually cultured on NGM plates at room temperature (22°C-23°C) unless specified otherwise. P0 animals with more than 100 post-injection progeny and at least 20 Rollers were selected— except at 100 ng/µl and 200 ng/µl of dsDNA donor where number of Rollers can be lower than 20 due to toxicity— and their F1 progeny were scored between 72 and 90 hours post-injection. All the F1 progeny from each brood were mounted onto 2% agarose pads and screened under fluorescence microscope for GFP or mCherry expression. GraphPad Prism (Version 8.4) was used to perform statistical tests and calculate P-values.

### Oligos and donors

End-modified donors were generated by PCR using oligos with 5′ SP9 modifications (IDT). Oligos used for to generate *hrde-1* and *F53H1*.*1 gfp* donors also contain 15bp linkers on either end of *gfp* which also serve as PCR primers. Sequences of all the crRNAs and oligos are provided in Supplemental Material, **Table S2** and **Table S3** respectively.

### Data availability

The authors state that all data necessary for confirming the conclusions presented in the manuscript are represented fully within the manuscript. All the reagents are available upon request.

## RESULTS

### Melting the donor dramatically stimulates HDR for longer edits

We recently showed that melting and reannealing donor molecules to create asymmetric donors with single-stranded homology arms can improve the frequency of CRISPR-mediated homology-directed insertions among transformants that were positive for a transformation marker (Dokshin *et al*. 2018). Because transformation markers can cause confounding effects or toxicity, we decided to conduct an initial study in which markers were omitted altogether. For this purpose, we chose to target the insertion of *gfp* into the easily scored *glh-1* locus, which encodes a VASA-related DEAD-box protein that localizes robustly to germline perinuclear foci known as P granules or nuage.

We prepared the *gfp* donor by PCR using primers tailed with 35 nt of homology to the *glh-1* locus (**Figure 1A**). In order to separately analyze the consequences of melting and of generating single-stranded overhangs we prepared three types of donor, (i) PCR products that were never melted, “unmelted donors,” (ii) “melted donors” that were heated and allowed to reanneal, and (iii) “asymmetric melted donors” that were prepared by heating a mixture of two overlapping *gfp* PCR products (one with 35-bp homology to *glh-1* at each terminus and one without (Dokshin *et al*. 2018). For simplicity, we refer to denaturing and quickly cooling the donor as “melting,” (see Methods). We injected each type of donor along with Cas9-guide-RNPs targeting *glh-1* into the core cytoplasm of the pachytene syncytium just distal to the gonad turn. Ideal injections result when the flow of the injection solution extends bilaterally from the injection site into the queue of oocytes at the proximal end and into mitotic region at the distal end (Mello *et al*. 1991). Only animals with two such injections—one per arm—were analyzed.

**Figure 1.**
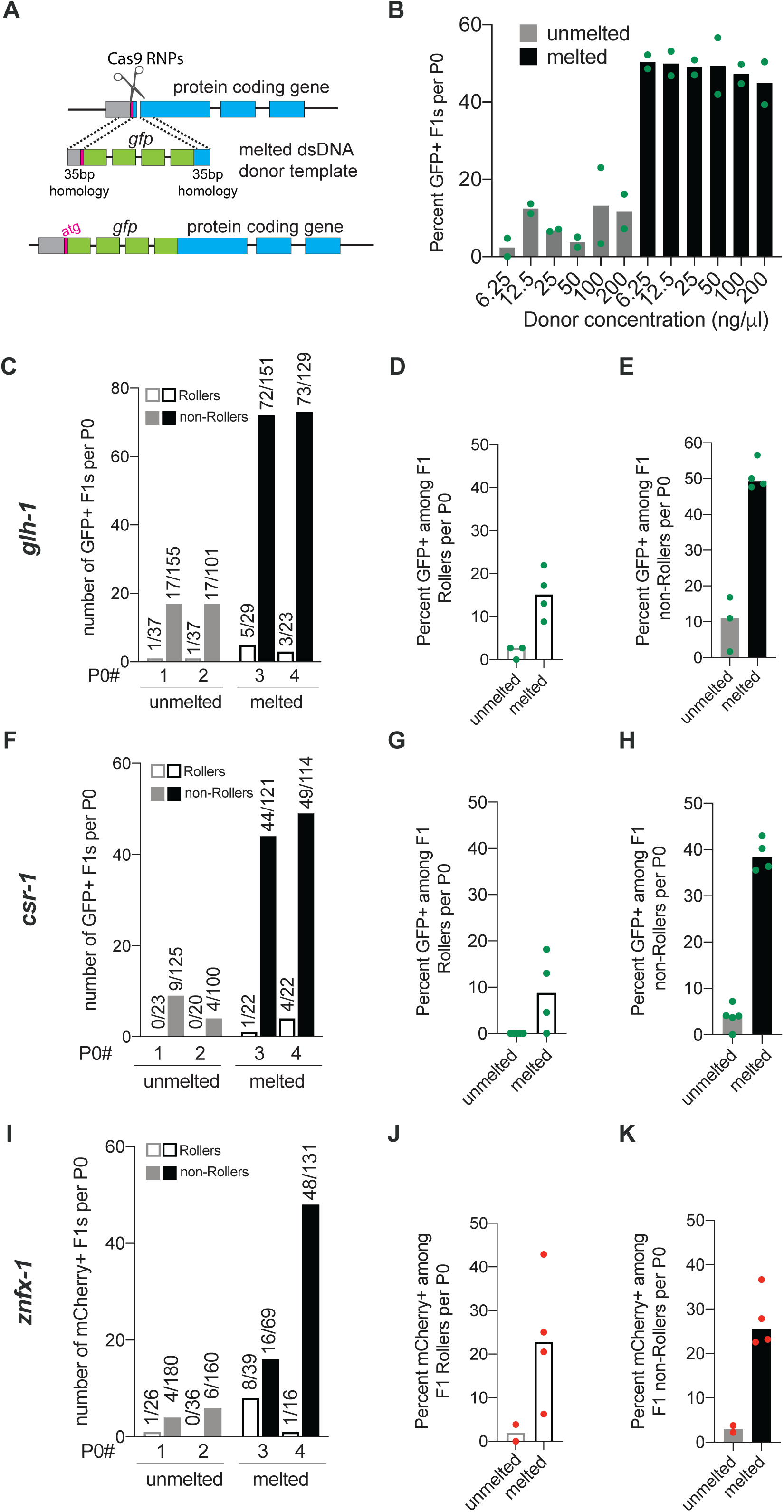
Melting dsDNA donors potentiates homology directed repair in *C. elegans*. **A**) Schematic representation to insert *gfp* at the N-terminus of a protein coding gene immediately down stream of start codon (atg) using symmetric melted dsDNA donors and Cas9-guideRNA ribonucleoproteins (RNPs) is shown; grey segment represents sequence upstream of the start codon. Precise repair (HDR) enables fluorescent protein expression. (**B**) HDR efficiencies at the *glh-1* locus using symmetric unmelted (grey bars) or melted donors (black bars) with *rol-6* injection marker at indicated concentrations (n=2 broods) is plotted as percentage of F1s expressing GFP per injected animal (P0). Using unmelted and melted donors, HDR efficiencies at the *glh-1* locus is plotted as (**C**) number of fluorescence+ animals among Rollers and non-Rollers from two representative broods. Percentage of animals expressing fluorescence among, (**D)** Rollers and (**E**) non-Rollers, is plotted as percentage (n = 3 or 4 broods) for *glh-1* locus. Similarly, improvement in fluorescent protein insertion efficiencies with melted donors are shown for (**F-H**) *csr-1* and (**I-K**) *znfx-1* loci. Each data point represents the percentage of animals expressing fluorescent protein among F1s scored in each cohort per brood. Bars represent median. Number of fluorescence+ animals over number of animals scored is shown above the bars. Green dots represent GFP and red dots represent mCherry insertions.

As previously shown (Dokshin *et al*. 2018) the asymmetric melted donor outperformed the unmelted donor. The asymmetric *gfp::glh-1* donor yielded 381 GFP-positive transformants among 900 F1 progeny, or 42% of total post injection progeny. The unmelted symmetric donor in contrast yielded half as many edits, 161 GFP-positive transformants among 740 post-injection progeny, (22%). Surprisingly, the symmetric melted donor was just as effective as the asymmetric melted donor, yielding 331 GFP positives among 906 F1 progeny, (37%). Thus, when the entire brood is scored a melted symmetric donor was as effective as its asymmetric counterpart. For melted donors, the number of GFP positive edits equaled approximately two-fifths of all post injection progeny exceeding the total number of Roller transgenics typically recovered per injected animal (**Figure S1 and see below**).

### Efficient HDR occurs over a broad range of donor concentrations

To explore how the frequency of *gfp* edits varied over a range of donor concentration, we injected unmelted or melted *gfp::glh-1* donor at concentrations of 6.25 ng/µl, 12.5 ng/µl, 25 ng/µl, 50 ng/µl, 100 ng/µl and 200 ng/µl (25 ng = 0.04 pmol). In order to control for injection quality, each injection mix included 40 ng/µl of the *rol-6(su1006)* co-transformation marker. For each donor mix, we injected 5 to 7 worms, singled those receiving optimal bilateral injections, and further analyzed two worms that made at least 100 post-injection progeny, including at least 20 Rollers. We then screened all the post-injection progeny—Roller and non-Roller—for germline GFP expression. We noted that the overall percentage of *gfp* insertions per injected animal (40%–50% for melted donors) (**Figure 1B**) was similar to levels achieved when the *rol-6* marker was omitted (**Figure S1**), suggesting that the *rol-6* marker does not interfere with the overall efficiency of editing. Surprisingly, the frequency of GFP-positive progeny per injected animal remained similar over a 32-fold range of donor concentrations. Melted donors consistently outperformed unmelted donors at every concentration (**Figure 1B**). These results suggest that, even at the donor concentration of 6.25 ng/µl, the HDR efficiency may be near saturation. At donor concentrations above 25 ng/µl, the frequency of Rollers per injected animal declined, suggesting that these higher concentrations cause toxicity (**Figure S2**). Taken together, these findings suggest that melted donors provide high rates of HDR with low toxicity over donor concentrations in the range of 6.25 ng/µl (0.01 pmol/µl) to 25 ng/µl (0.04 pmol/µl). Based on these findings we chose to use 25 ng/µl of donor in further investigations.

We next wished to examine how editing efficiencies vary among the Roller and non-Roller cohorts of post-injection progeny. We found that melted donors out-performed unmelted donors in both Roller and non-Roller cohorts (**Figure 1, C-E**), yielding several dozen *gfp* edited progeny per injected animal (as shown in two representative broods, **Figure 1C**). Strikingly, the fraction of GFP expressing progeny was much higher among non-Rollers (49%) (**Figure 1E**) compared to Rollers (15%) (**Figure 1D**).

To confirm the generality of these findings, we targeted two additional germline-expressed genes, *csr-1* and *znfx-1* (**Figure 1, F-K**). In both cases, melted donors consistently outperformed unmelted donors for *gfp* and *mCherry* insertions respectively (**Figure 1, F and I)**. When melted donors were used, the fraction of animals with precision insertions was approximately ∼10-fold higher than levels obtained with unmelted donors. This enhancement was observed in both the Roller (**Figure 1, G and J**) and non-Roller cohorts (**Figure 1, H and K)**. We also explored whether melted donors were beneficial for editing with Cas12a (CPF1) (EBBING *et al*. 2017) RNPs — which recognize an AT rich TTTV protospacer adjacent motif (PAM) sequence. Indeed, Cas12a editing yielded high HDR efficiencies comparable to those achieved with Cas9 RNPs for *gfp* insertion at both the *glh-1* and *F53H1*.*1* loci (**Figure S3**) (See **File S1** for protocol).

### Editing efficiency peaks later in the brood after the Roller cohort of progeny are produced

The finding that HDR events are more prevalent among non-Roller progeny might reflect different developmental competencies of germ nuclei to form these distinct types of transgenics. For example, distal pachytene germ nuclei may be more receptive to recombination between the target chromosomal locus and the *gfp* donor, whereas more proximal germ nuclei may be more receptive to forming extra-chromosomal transgenes driven by recombination between co-injected DNA molecules (see Discussion) (Mello *et al*. 1991). To examine these possibilities, we followed the production of Roller and GFP-positive progeny over the entire post-injection brood. Worms receiving ‘ideal’ bilateral injections of an editing mix prepared with melted *gfp*::*glh-1* donor (25 ng/µl) and *rol-6* co-injection marker (40 ng/µl) were cultured in two groups of 4 injected animals. Each group of animals was transferred every 4 hours to fresh plates to divide their broods into 12 segments over the next two days. Animals were transferred one more time on the third day (64 hours post-injection) thus dividing the progeny into 14 groups (**Figure 2A**). We then scored the frequency of Roller progeny and GFP-positive progeny in each segment.

**Figure 2.**
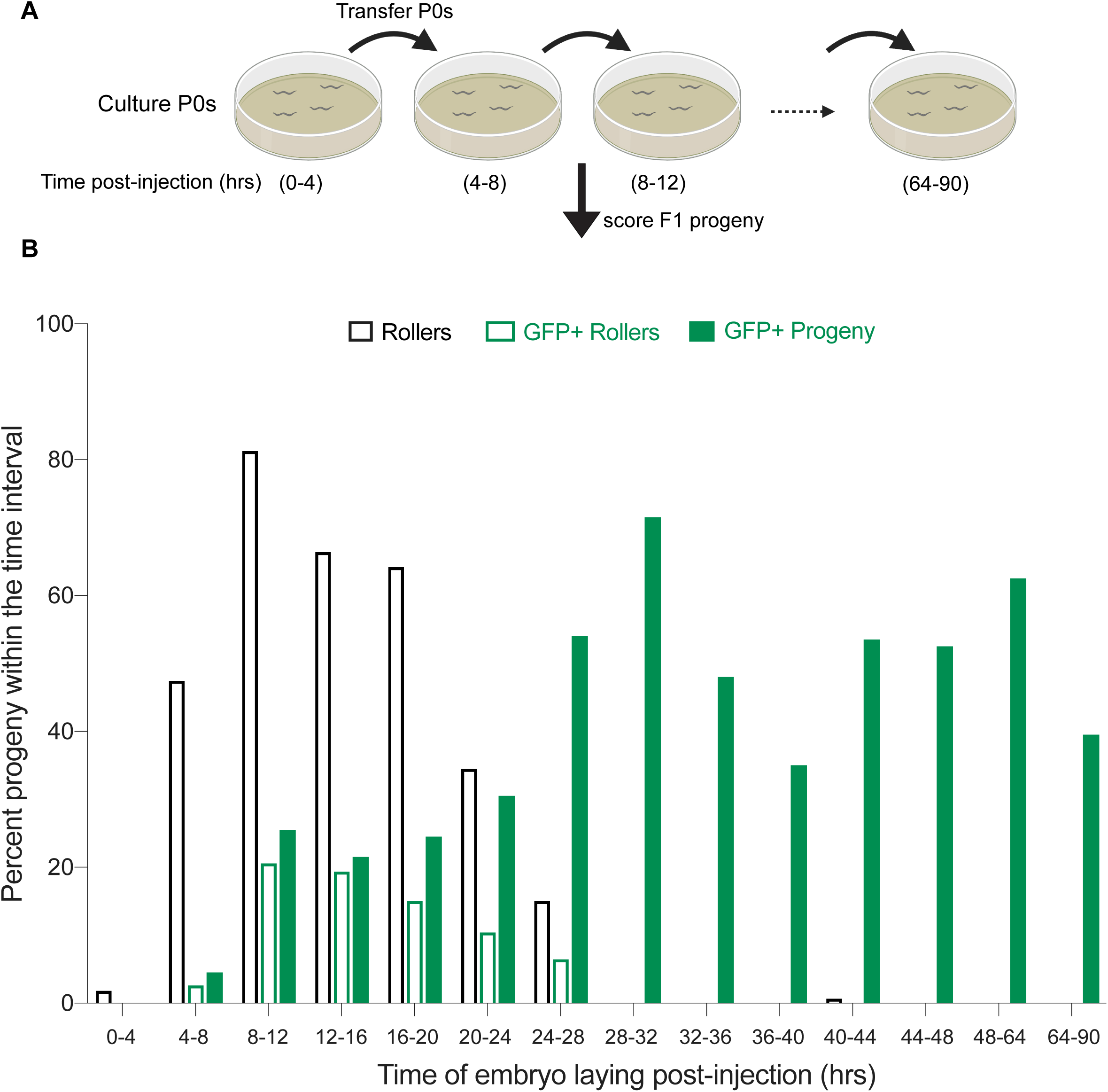
Editing occurs later in the brood after roller cohort. (**A**) Schematic representation of the experiment is shown. 4 Injected animals placed on a single plate were moved at indicated post injection time points and F1 embryos laid during the time-intervals were scored for GFP as adults. **(B)** Fraction of the progeny produced in each time window that are Rollers (open black bars), GFP+ Rollers (open green bars) and GFP+ progeny (Rollers and non-Rollers, solid green bars) are plotted as percentage. Bars represent mean value of two replicates and each replicate consists of 4 P0 animals injected with 25 ng/µl of symmetric melted donors and 40 ng/µl of *rol-6* co-injection marker. Animals were cultured at 18°C-20°C.

Consistent with the idea that Roller extra-chromosomal transgenes assemble in more proximal germ cells, nearly 100% of the Roller progeny were produced within the first 28 hours post injection. The frequency of Rollers peaked between 8 and 12 hours post-injection where Rollers comprised 81% of the 47 progeny produced in the interval. The frequency of Roller progeny remained above 60% until 20 hours post injection, declining to ∼30% then 13% over the next two 4-hour intervals. Rollers were virtually absent among progeny produced after 28 hours (**Figure 2B**). In striking contrast, the frequency of precision editing events was low within the first 24 hours and then appeared to plateau and remain high during the entire remainder of the brood (**Figure 2B**). For example, only 20% of the 306 progeny produced in the first 24 hours were GFP positive while an average of 54% were positive among the progeny produced thereafter (n=1327). Importantly, while GFP precision editing was less frequent within the first 24 hours (where Roller transgenics were found), precision editing was not under-represented within the Roller cohort. For example, we found that 24% of Rollers vs 20% of all animals produced in the first 24 hours were GFP positive (**Figure 2B**). Moreover, among GFP positive animals produced in this interval 60% were Rollers. Thus, the Roller marker positively correlates with *gfp* editing but does so within a cohort of progeny that precedes the optimal editing window for *gfp* insertion (See Discussion).

### Donor purity is crucial for best HDR efficiencies

Although *rol-6* transformation precedes the optimal window of *gfp* insertion (as shown above), we nevertheless found that the *rol-6* marker provides a valuable troubleshooting metric (**Figure S2**). For example, while attempting to knock-in *gfp* at two different loci (*hrde-1* and *F53H1*.*1*), *gfp* insertions were unexpectedly rare. These experiments were conducted using melted TEG-modified donors (Ghanta *et al*. 2018), which typically yield as many as 100 GFP+ progeny per injected worm. However, despite ideal injections that produced high numbers of Roller progeny, only 2 (average) Rollers were GFP positive per brood (spin-column, **Figure 3, A** and **D**). Scoring entire broods for GFP, we only obtained a maximum of 18 (*hrde-1*) (**Figure 3A**, P0# 2) and 13 (*F53H1*.*1*) (**Figure 3D**, P0#s 1 and 2) GFP-positive progeny per injected worm. The fraction of Rollers (spin-column, **Figure 3, B and E)** or non-Rollers (spin-column, **Figure 3, C and F)** expressing GFP stayed below 8% at both the loci. Because the number of Rollers per injected animal was near the optimal range, we reasoned that the injection quality was good, injected animals were healthy, and the injection mixture was non-toxic.

**Figure 3.**
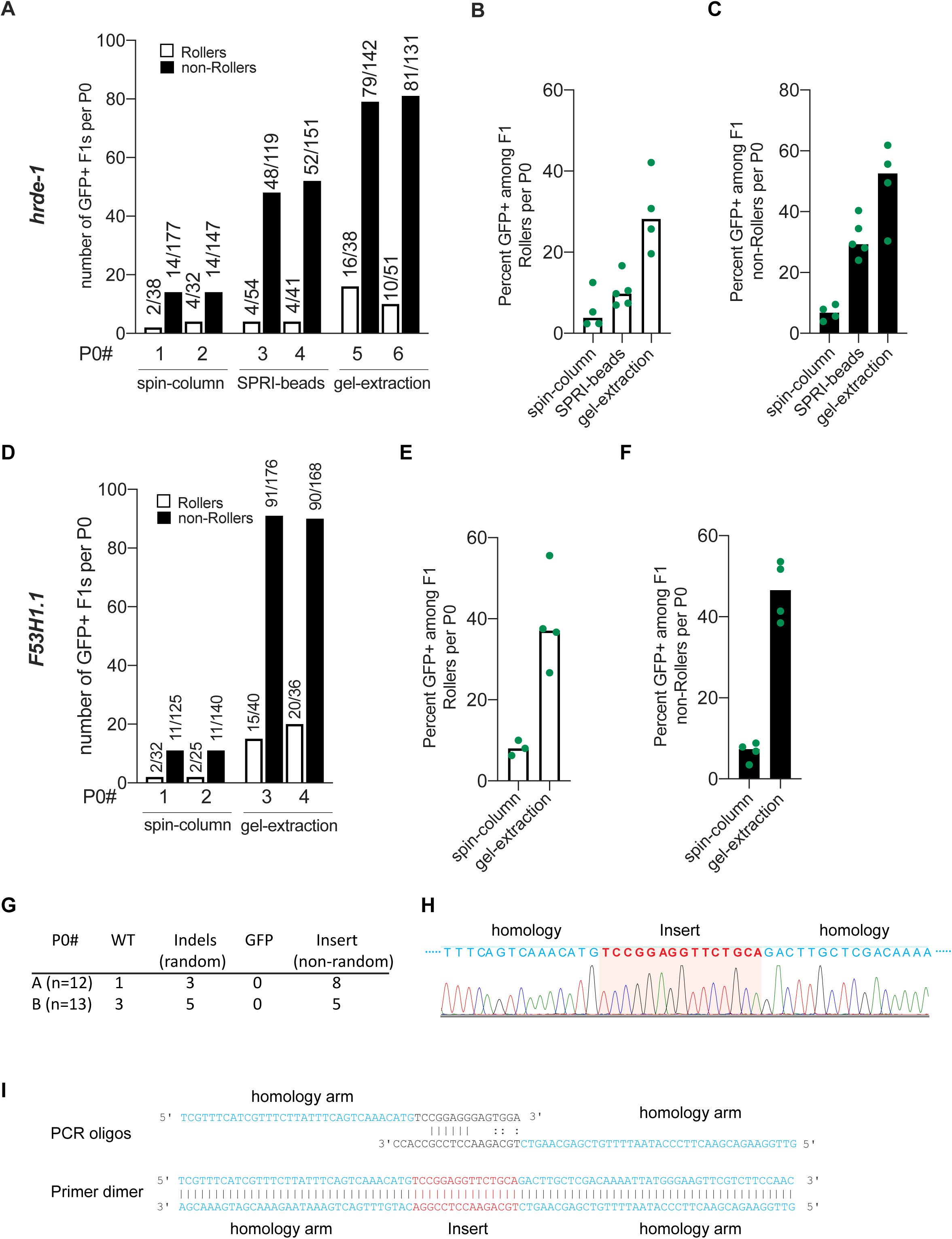
Purity of donor DNA is crucial for best HDR efficacy. HDR efficiencies of donors prepared by different methods of purification are plotted for *hrde-1* and *F53H1*.*1*. (**A**) Number of GFP+ F1 Rollers and non-Rollers from two representative broods are plotted. GFP+ animals among, (**B**) Rollers and (**C**) non-Rollers is plotted as percentage of animals scored in each cohort per brood. Similarly, (**D-F**) HDR efficiencies are plotted for *F53H1*.*1* locus. (**G**) Insertions and deletions identified at *hrde-1* target site in F1 Rollers from two P0s (spin-column), (**H**) sanger sequencing trace of the 15bp non-random insert for a homozygous F2 animal. Partial homology arms of the donor are shown in blue and the sequence that got inserted into the genome is shown in red. (**I**) Schematic representation of predicted primer dimer formation is shown with 6bp perfect match and mismatched 3′ tails. Part of each oligo that is homologous to the PCR template plasmid is shown in black (linkers on either end of *gfp*) and the homology arms are shown in blue and the sequence (Insert) that would get inserted through HDR is shown in red. All the donors were 5′ TEG-modified and melted. Gel-purified donors were further cleaned-up with SPRI beads (See **File S1**).

To understand why editing was so infrequent, we randomly sequenced the target site in 25 F1 Rollers. In 21 of 24 Rollers, we identified non-wildtype sequences at the target site (**Figure 3G**), indicating that double-strand breaks were not the limitation. Importantly, none of these 21 Rollers contained *gfp* insertions (**Figure 3G**). Upon reading the sequencing trace, we found that 13 F1 animals contained a 15-bp insertion precisely where GFP sequences should have inserted (**Figure 3G and 3H**). To our surprise, this short sequence perfectly matched a segment of the PCR oligo sequences (**Figure 3I**), and thus could be explained by insertion of a primer fragment or primer-dimer that was produced inadvertently during donor preparation. To test this possibility, we purified the *gfp* donors by size-exclusion using SPRI paramagnetic beads or by gel-extraction. Purifying the *hrde-1* donor with SPRI beads (optimized to exclude fragments smaller than 300 bp) modestly increased the percentage of GFP-positive progeny to 10% of F1 Rollers (n=212; **Figure 3B**) and 32% of non-Roller progeny (n=625; **Figure 3C**). By contrast, gel-purified *hrde-1* donor dramatically increased the percentage of GFP-positive progeny to 29% of F1 Rollers (n=163; **Figure 3B**) and 49% of non-Rollers (n=538; **Figure 3C**), with as many as 95 GFP-positive progeny from one injected worm (**Figure 3A**, P0#5). Similar results were obtained after gel purification of the *F53H1*.*1* donor (**Figure 3, D-F**). These findings demonstrate the utility of the Roller marker as a metric for troubleshooting the editing protocol and reveal the importance of removing PCR-based contaminants from donor preparations to achieve best knock-in efficiencies.

## DISCUSSION

We initiated these investigations to explore why long (∼1-kb) DNA donors were less efficacious than short single-stranded oligonucleotide (ssODN) donors in *C. elegans*. We have shown that melting the donor DNA dramatically enhances precision editing, enabling efficient editing with shorter homology arms and at significantly lower donor DNA concentrations than previously recommended (Paix *et al*. 2015; Paix *et al*. 2017; Dokshin *et al*. 2018). We show that as many as 100 precisely edited progeny can be obtained from a single injected animal, an editing efficiency of nearly 50% of post injection progeny, and far exceeding the typical frequency of progeny transformed with simple extrachromosomal arrays (**Figure S2**) (Mello *et al*. 1991).

Importantly, whereas the production of Roller transgenic progeny peaks during the first 24 hours post-injection, *gfp* edits peak after 24 hours and remain high through the remainder of the injected animals brood. Previous studies also reported that most *gfp*-edited animals are produced on the second day after injection (Paix *et al*. 2014). These findings suggest that developmental differences between distal (less mature) and proximal (more mature) germ nuclei may favor formation or acquisition of distinct transgene types. For example, perhaps the large *rol-6* plasmid molecules are excluded from germ nuclei, and instead rapidly assemble into cytoplasmic extrachromosomal arrays that are swept by the germ plasm into developing oocytes, and only enter nuclei after fertilization (as previously suggested (Mello *et al*. 1991)). A size limitation on nuclear uptake may explain why we and others have found that donors over 2 kilo-basepairs yield few editing events (unpublished results) (Paix *et al*. 2016; Farboud *et al*. 2019).

The observation that *gfp* editing peaks later, approximately 28 hours post injection, and then remains high, suggests either that proximal germ nuclei tend to exclude the donor, or that more distal nuclei are more receptive to recombination. Based on an ovulation rate of 23 minutes (Mccarter *et al*. 1999), the appearance and persistence of GFP-positive progeny is consistent with editing in nuclei that were in pachytene (i.e., undergoing meiotic recombination) at the time of injection. Whatever the reason for the HDR enhancement caused by melting the donor it is striking that the extrapolated rates of precision *gfp* insertion within these pachytene nuclei range as high as 70%.

Donor purity is crucial to achieve high knock-in efficiencies of long inserts. Contaminating primer dimers that contain homology arms can compromise HDR efficiency by integrating at the target site. Removing these contaminants by gel-extracting the donors dramatically increased *gfp* knock-in efficiencies. Similarly, as a time saving alternative to gel-extraction we found that purification using SPRI paramagnetic beads also improves HDR efficiencies, however using the optimal ratio of beads to PCR reaction was critical to removing the shorter contaminants (See protocol **File S1**).

We do not know why melting the donor stimulates HDR. We obtained similar HDR rates across the entire range of donor concentrations, indicating that donor concentrations were saturated (or nearly so) at the lowest dose tested. Yet, melting the donor increased the HDR rate several fold at each concentration. Thus, melting stimulates recombination by acting on events or mechanisms that are independent of donor concentration. Conceivably, melting induces structural changes— e.g., denaturation bubbles caused by incomplete reannealing—that promote active nuclear uptake or directly stimulate repair. For example, single-stranded regions from incomplete re-annealing could promote strand invasion or act as damage signals that recruit trans-acting factors that facilitate HDR. Indeed, preliminary studies suggest that when we slow the cooling step to promote re-annealing, melted donors perform about half as well (**Figure S4**). Interestingly, melting did not stimulate the already high HDR efficiency of a shorter 400 nt donor (Data not shown). Clearly more work is needed to fully explore and understand how donor-melting promotes HDR efficiency.

Undoubtedly, the high efficiencies of precision editing achieved here owe both to the easy access of worm pachytene germ cells to microinjection and to the remarkable receptiveness of these cells to HDR. A parallel study suggests that editing is enhanced even further when donor 5′ ends are modified with tri-ethylene glycol (TEG) (Ghanta *et al*. 2018). Importantly, the combination of melting and TEG modifications increases the proportion of *gfp*-sized edits among the easily identified Roller progeny cohort by approximately twenty-fold from 1-2% to 20-40%. For experienced injectors, a single optimally injected animal can yield more than 100 GFP knock-ins (nearly two thirds of post-injection progeny), dramatically enhancing the ease and efficiency of genome editing. Given these high HDR efficiencies even researchers with little worm experience can now readily adopt this facile genetic animal model.

## Acknowledgements

This work was funded by Howard Hughes Medical Institute (C.C.M.) and NIH R37 GM058800 (C.C.M). C.C.M is a Howard Hughes Medical Investigator. We thank Daniel Durning for technical assistance with preparation of *mCherry::znfx-1* injection mixes, Takao Ishidate for providing plasmid template for *linker-gfp-linker* donor PCRs and Darryl Conte, Jr. for critical review of the manuscript.

## Competing interests

C.C.M is a co-founder and Scientific Advisory Board member of CRISPR Therapeutics. Some of the findings described here are part of the patent applications filed by the University of Massachusetts Medical School on which the authors are inventors.

## Supplementary Figures

**Figure S1:**
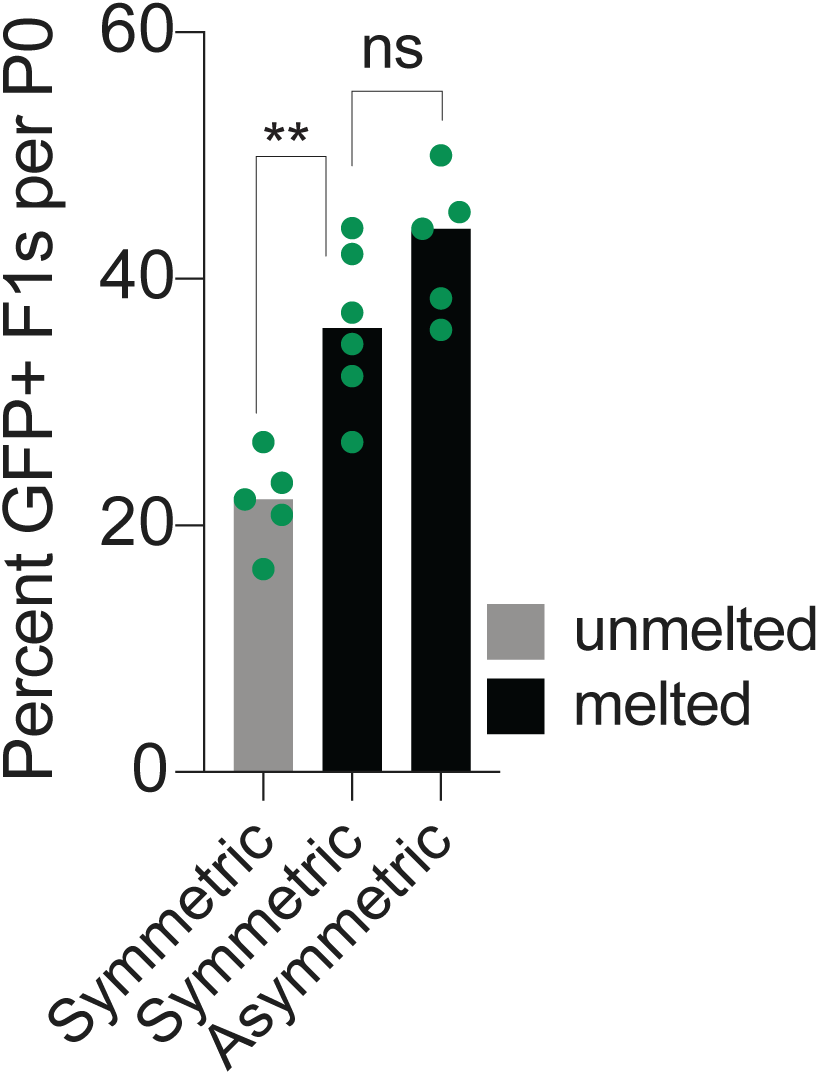
Melted dsDNA donors promote homology directed repair. HDR efficiencies at the *glh-1* locus using symmetric (unmelted or melted) and asymmetric donors (n=5 or 6 broods) without *rol-6* injection marker. Each data point (green) represents the percentage of animals expressing GFP among F1s scored per brood. Bars represent median.P-values (**, 0.0087 and ns, 0.1255) were determined by Mann Whitney test (unpaired, non-parametric)

**Figure S2.**
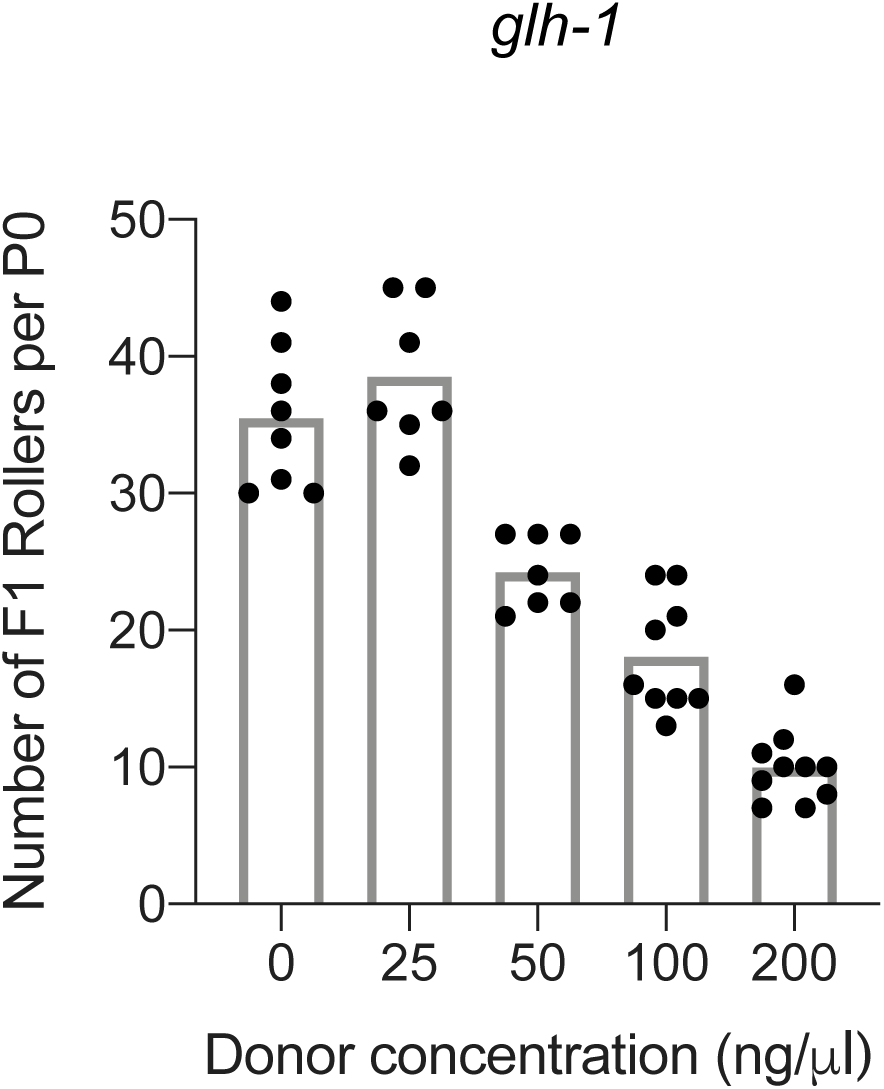
High donor concentrations are toxic and reduce the number of Rollers. Injection mixes contain Prf4::*rol-6(su1006)*, Cas9 protein, crRNA targetting *glh-1* locus and *gfp::glh-1* dsDNA donor with ∼35bp homology arms at indicated doses. Each dot represents the number of F1 rollers obtained per P0 animal and the bar represents mean; (n=7 to 10 broods per condition). dsDNA donors were not melted.

**Figure S3.**
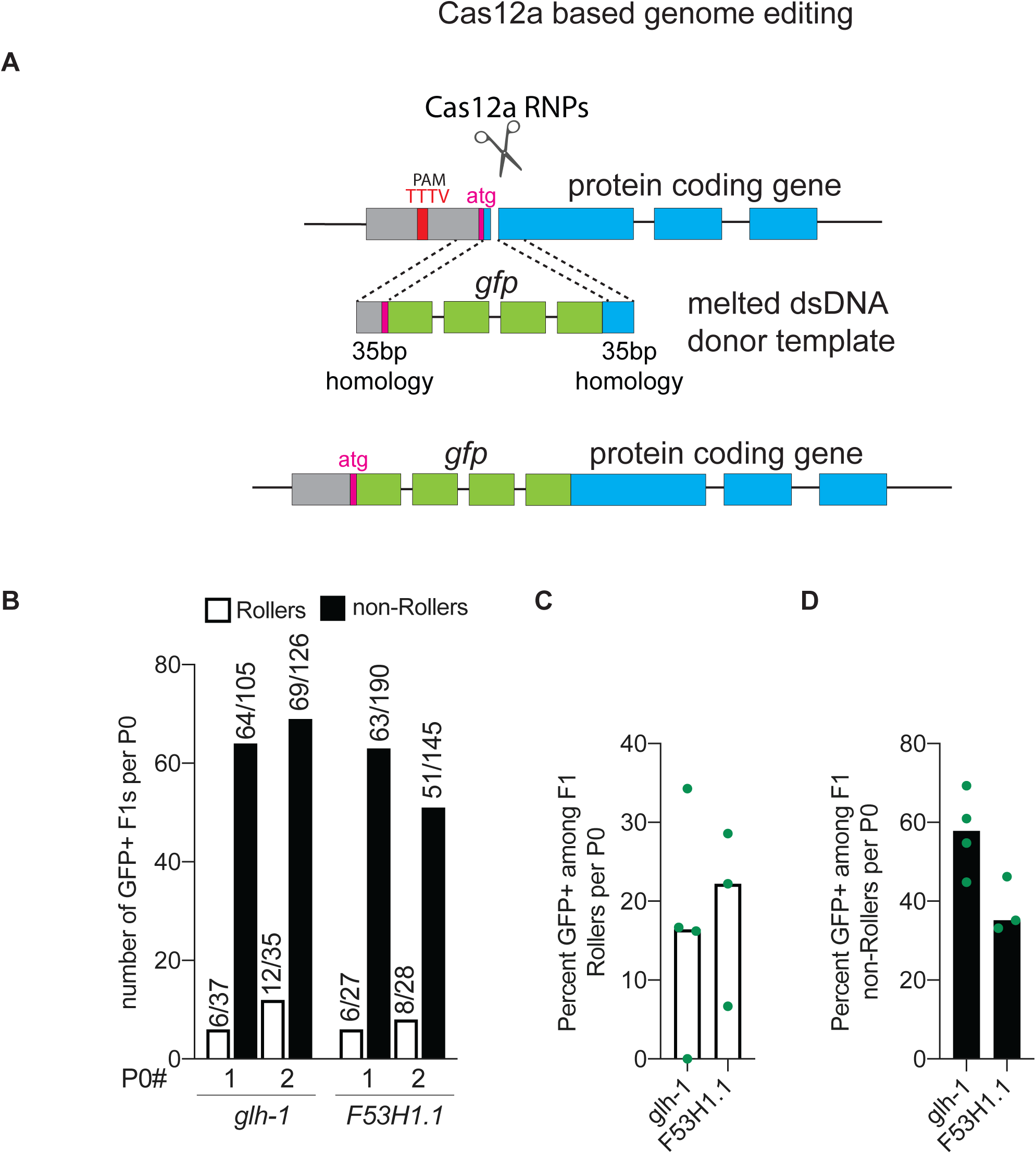
Precise genome editing using Cas12a nuclease and melted dsDNA donors. (**A**) Schematic represention of template dependent editing with Cas12a and melted dsDNA donor to insert *gfp* at the start codon (atg). Protospacer adjacent motif (PAM) for Cas12a system is TTTV, where V is A, C or G. HDR efficiencies at *glh-1* and *F53H1*.*1* loci are plotted as (**B**) number of GFP+ F1 animals among two representative broods, (**C**) percentage of GFP+ animals among F1 Rollers and (**D**) percentage of GFP+ animals among F1 non-Rollers; n= 3 or 4 broods. Number of GFP+ animals over number of animals scored are shown above the bars. Each data point represents the percentage of animals that are GFP+ among F1s scored in each cohort per brood and bars represent the median.

**Figure S4.**
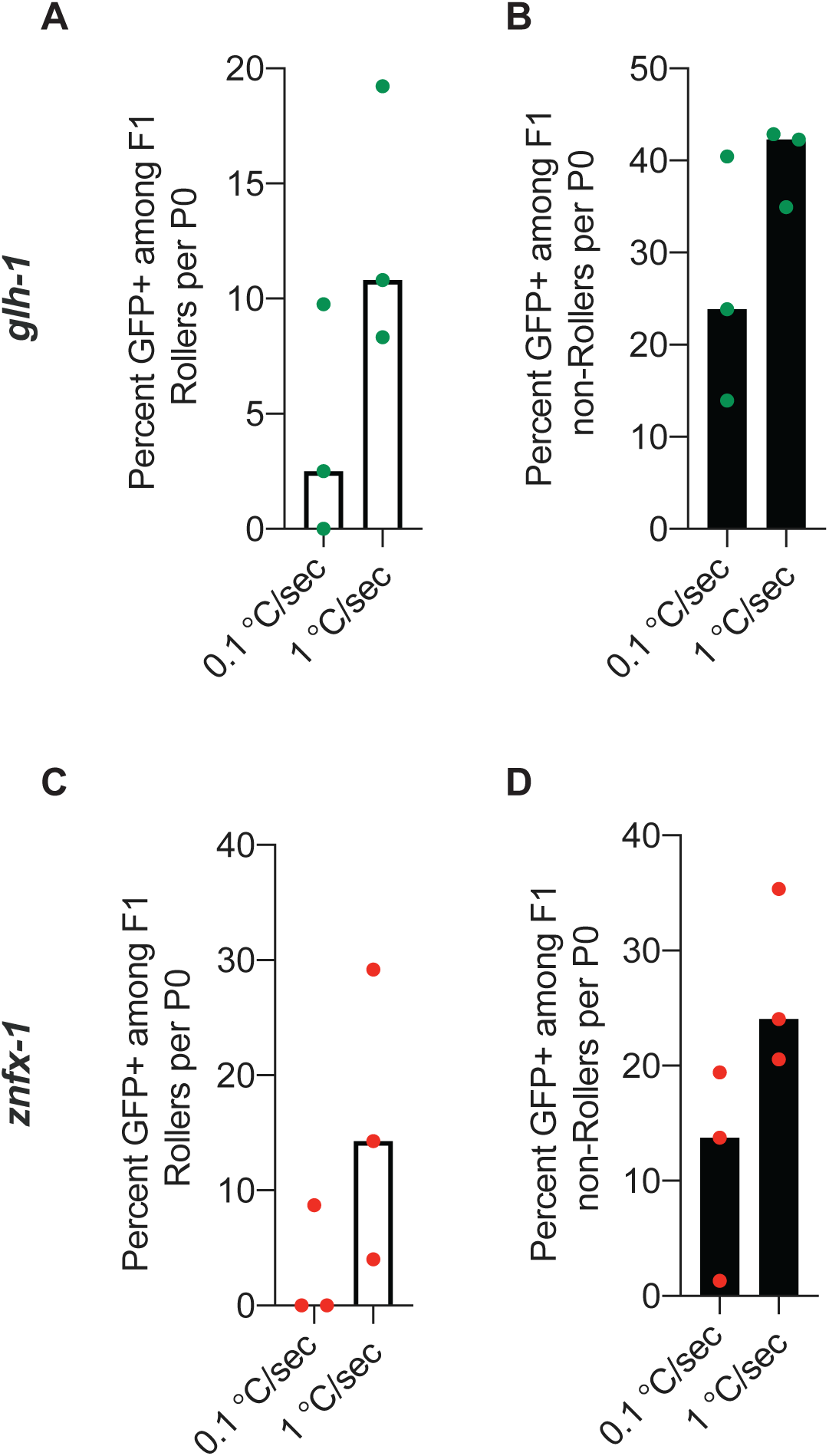
Quickly cooled donors act as better repair templates than slowly cooled donors. *gfp* (green dots) insertion efficiencies at *glh-1* loci are plotted for (**A**) Rollers and (**B**) non-Rollers using slow (0.1 °C/sec) and quick (1 °C/sec) cooled donors as percentage. (**C and D**) *mCherry* (red dots) insertion efficiencies at *znfx-1* locus. Each data point represents an F1 brood and bars represent median. Thermal cycler program for slow cooling: 95 °C - 2:00 min; 85 °C - 1:00 min; 75 °C - 1:00 min, 65 °C - 1:00 min, 55 °C - 1:00 min, 45 °C - 1:00 min, 35 °C - 1:00 min, 25 °C - 1:00 min, 4°C- hold. Ramp down rate: 0.1 °C/sec. See **File S1** for quick cooling conditions.

## Supplemental Information

1. Figure S1- Melted dsDNA donors promote homology directed repair

2. Figure S2- High donor concentrations are toxic and reduce the number of Rollers

3. Figure S3- Precise genome editing using Cas12a nuclease and melted dsDNA donors

4. Figure S4- Quickly cooled donors are better repair templates than slowly cooled donors

5. Table S1- List of *C. elegans* strains

6. Table S2- Sequences of crRNAs

7. Table S3- Sequences of Oligos

8. File S1- Detailed editing protocol

**Supplementary Table S1:** List of *C. elegans* strains.

| Genotype | Strain Name |
| --- | --- |
| F53H1.1(ne4815[gfp::F53H1.1]) IV | WM703 |
| glh-1(ne4816[gfp::glh-1]) I | WM704 |
| csr-1(ne4483[flagx3::linker::tev] IV | WM705 |
| znfx-1(ne4817[mcherry::znfx-1]) II | WM706 |
| hrde-1(ne4818[gfp::hrde-1]) III | WM707 |
| csr-1(ne4819[flagx3::linker::gfp::tev::csr-1]) IV | WM708 |

**Supplementary Table S2:**
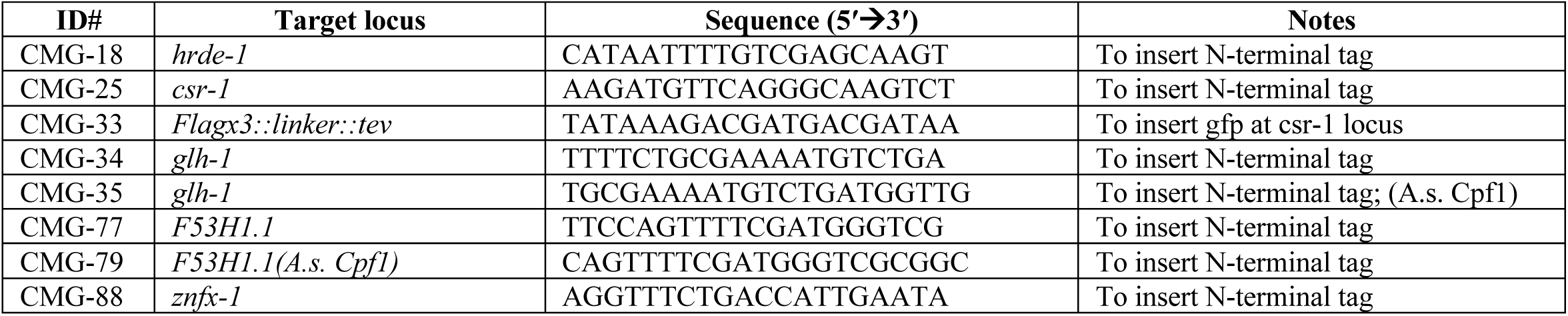
Sequences of crRNAs.

**Supplementary Table S3:**
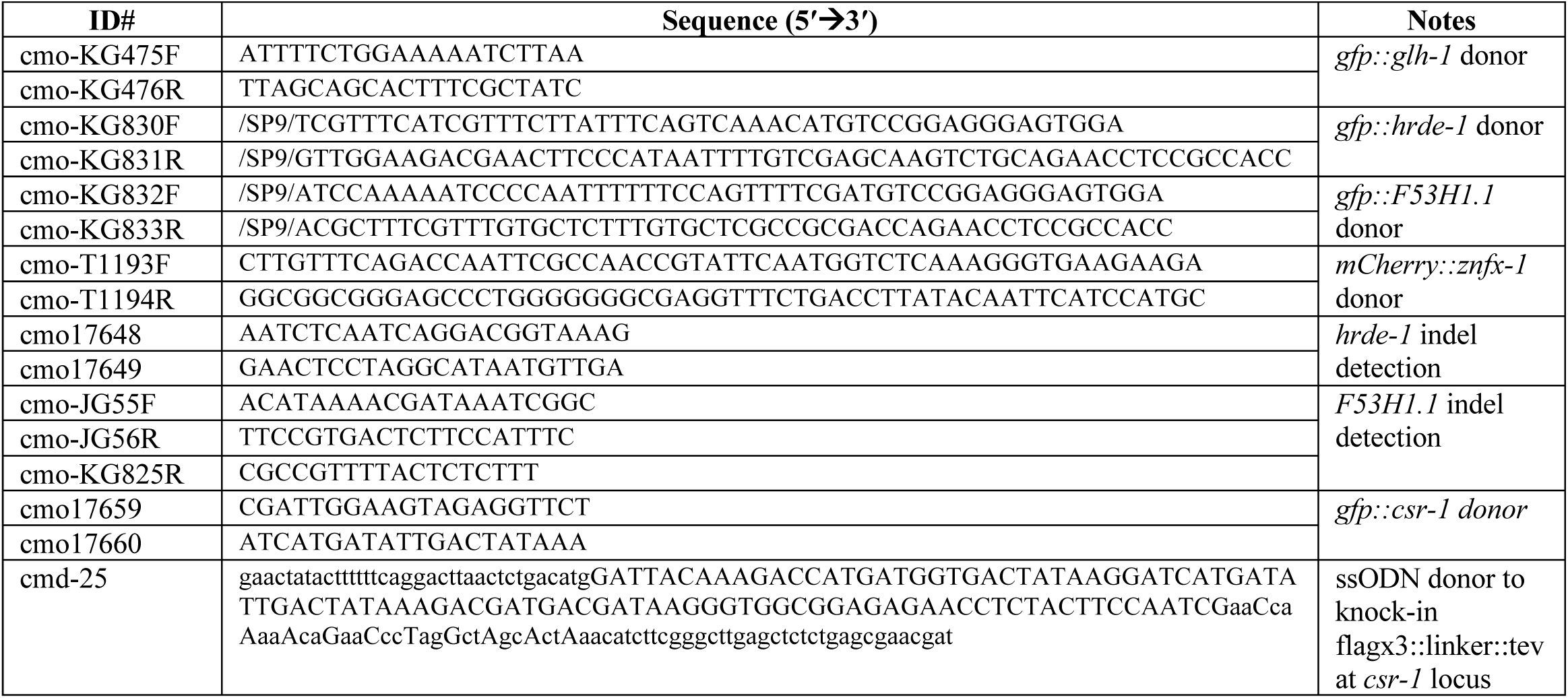
Sequences of oligos.

## Cas9 Based Genome Editing

### I. Materials

1. *S. pyogenes* Cas9 3NLS (10 µg/µl, IDT)
2. tracrRNA (IDT)
3. crRNA (2 nmol or 10 nmol, IDT)
4. ssODN 4 nmol Ultramer (standard desalting, IDT)
5. PRF4::*rol-6(su1006)* plasmid (high quality Midi or Maxiprep)
6. SPRI paramagnetic beads (AMPure XP, Beckman Coulter)

#### Re-suspension (Stock Solutions)

1. Aliquot 0.5 µl (5 µg or 30 pmol) of Cas9 protein and store at -80°C (avoid freeze/thaw cycles)
2. tracrRNA – 0.4 µg/µl (18µM) in IDT nuclease free duplex buffer, store at -20°C (aliquots at -80°C)
3. crRNA – 0.4 µg/µl (34 µM) in TE PH 7.5 (IDT), store at -20°C (aliquots at -80°C)
4. ssDNA oligo donor – 1 µg/µl in ddH2O, store at -20°C
5. PRF4::*rol-6 (su1006):* 500 ng/µl, store at -20°C

### II. Injection mixture preparation

Add components of the injection mixture to the tube containing Cas9 in the following sequence:

1. Cas9 – 0.5 µl of 10 µg/µl stock (30 pmol)
2. Add tracrRNA – 5 µl of 0.4 µg/µl stock (90 pmol)
3. Add crRNA – 2.8 µl of 0.4 µg/µl stock (95 pmol) (if two guides are needed add 1.4 µl of each)
4. Pipette the mixture gently several times and incubate @37°C for 15 minutes. In our experience adding any double stranded DNA before RNP complex formation reduces HDR efficiency.
5. Add ssODN donor – 2.2 µl of 1 µg/µl stock (or)
Add melted dsDNA – 500 ng (final concentration: 25 ng/µl for ∼1kb donors or 45 fmol/µl)
6. Add PRF4::*rol-6 (su1006)* plasmid – 1.6 µl of 500 ng/µl stock
7. Add nuclease free water to bring the final volume to 20 µl and pipette gently several times.
8. To avoid needle clogging, centrifuge the mixture @14000rpm for 2 min, transfer about 17 µl of the mixture to a fresh tube and keep the tube on ice; proceed to loading the needles.

*Notes:*

- *All the above steps in section II can be performed at room temperature*
- *Aggregation is not an issue under these Cas9 concentrations*.
- *Final injection mixture can be stored at 4°C and re-used for several months (up to 6 months) without compromising efficiency; we have not yet tested mixes that are older than 6 months*.

## Cas12a (Cpf1) Based Genome Editing

### I. Materials

1. A.s. Cas12a Ultra (10 µg/µl, IDT)
2. Cpf1-crRNA 21 bases long (2 nmol or 10 nmol, IDT)
3. ssODN 4 nmol Ultramer
4. PRF4::*rol-6(su1006)* plasmid (high quality Midi or Maxiprep)
5. 7. SPRI paramagnetic beads (AMPure XP, Beckman Coulter)

#### Re-suspension (Stock Solutions)

1. Aliquot 0.5 µl (5 µg or 32 pmol) of Cas12a protein and store at -80°C (avoid freeze/thaw cycles)
2. Cas12a-crRNA – 40 µM in TE PH 7.5 (IDT), store at -20°C (aliquots at -80°C)
3. ssDNA oligo donor – 1 µg/µl in ddH2O, store at -20°C
4. PRF4::*rol-6 (su1006):* 500 ng/µl, store at -20°C

### II. Injection mixture preparation

Add components of the injection mixture to the tube containing Cas9 in the following sequence:

1. Cas12a – 0.5 µl of 10 µg/µl stock (32 pmol)
2. Add cas12a-crRNA – 2.5 µl of 40 µM stock (100pmol)
3. Add TE PH 7.5 – 3.0 µl
4. Pipette the mixture gently several times and incubate @37°C for 15 minutes
5. Add ssODN donor – 2.2 µl of 1 µg/µl stock (or)
Add melted dsDNA – 500 ng (final concentration: 25 ng/µl for ∼1kb donors or 45 fmol/µl)
6. Add PRF4::*rol-6 (su1006)* plasmid – 1.6 µl of 500 ng/µl stock
7. Add nuclease free water to bring the final volume to 20 µl and pipette gently several times.
8. To avoid needle clogging, centrifuge the mixture @14000rpm for 2 min, transfer about 17 µl of the mixture to a fresh tube and keep the tube on ice; proceed to loading the needles.

*Notes:*

- *All the above steps in section II can be performed at room temperature*
- *TE is added in step 3 for easier pipetting; by further diluting the crRNA stock this step can be omitted*.

## III. Donor Design and Generation ssODN donors

To generate ssODN donor, add 35 bases of 5′ homology sequence in front of the tag (or mutations) and 35 bases of the 3′ homology sequence at the end. Mutate the PAM site or the guide binding sequence if it is not already disrupted by the insert. If the guide binding sequence is mutated or if silent mutations are introduced between the guide cleavage site and the desired insertion site, length of homology sequence should be 35bp from the last mutation.

### dsDNA donors

Generate dsDNA donors by PCR either by using unmodified oligos or 5′ SP9-modified oligos.

1. Order unmodified (or 5′ SP9 modified) oligos with standard desalting (IDT); 35nt as homology arms and 20nt complementary to insert (eg: GFP). SP9 modifications are available at 100nmol scale from IDT.
2. Perform PCR with an insert-containing plasmid as the template for amplification; use High-Fidelity polymerase.
3. Run a few microliters of PCR on agarose gel to check if a single bright band is obtained. If non-specific amplification is observed, set up a temperature gradient and find the optimal temperature.
4. PCR clean-up: use one of the following three options depending on your experimental conditions.
  a. Purify the PCRs using spin-columns and elute DNA in 20 µl of nuclease free water. Generally, column purification is sufficient, and you may proceed to step 5. However, some primer pairs produce long (∼80bp) primer dimers that may contain the entire homology arms. Spin-columns may not be able to remove dimers of this length completely. We found that these short “dimer donors” are preferentially used as templates over full-length donors with the desired insert (such as GFP). *Note*: Dimers may or may not be visible on the agarose gel.
  b. If dimer formation is a concern, use 0.6x SPRI beads (AMPure XP) to perform the clean-up instead of spin-columns. For example: add 60 µl of beads to 100µl of PCR, wash with 70% ethanol twice, elute in nuclease free water (refer to the bead manufacturer’s protocol for further details).
  c. If primer dimers are clearly visible on the gel, then it is best to gel-extract the DNA. However, gel extracted DNA can be toxic, presumably due to the presence of guanidine hydrochloride (component of binding buffer) in the final elute. To reduce salt contamination, incubate the column with wash buffer for 10 min before centrifugation; repeat washes 2-3 times. Strong absorbance at 230nm on Nanodrop suggests GuHCl contamination. For best results, gel-extracted DNA should be further purified with 1x to 1.5x AMPure XP beads (strongly recommended).
5. After purification, dilute a portion of dsDNA PCR donor to 100 ng/µl and transfer about 5.5 µl to a PCR strip tube and proceed to the heating step.
6. Heat to 95 °C and cool to 4 °C using thermal cycler (95 °C-2:00 min; 85 °C-10 sec, 75 °C-10 sec, 65 °C-10 sec, 55 °C-1:00 min, 45 °C-30 sec, 35 °C-10 sec, 25 °C-10 sec, 4 °C-hold. Ramp down at 1 °C/sec at every step).
7. Add melted donor DNA to rest of the injection mixture only after pre-incubating RNP complexes.

*Note: we store purified donors at -20* °C and *melt them right before adding to the injection mix. We have not explored storage and re-use of melted donors*.

## IV Micro-injection and Screening

1. Inject 5 to 10 young adults and transfer them onto individual plates. If both arms of the hermaphrodite gonad are injected, a good injection should yield 20 to 40 F1 Rollers.
2. After about 72 hours post injection, score for number of F1 Rollers and choose 2 plates with the highest number of Rollers.
*Note:* We generally culture the injected animals at room temperature (∼22°C-23°C).
3. a. For ssODN-based editing: choose 2 P0 plates that segregate the highest number of F1 Rollers; pick about 24 F1 Rollers from these 2 plates and place them onto separate plates.
b. For dsDNA-based editing: Choose 2 plates that segregate the highest number of F1 Rollers and from these 2 plates, pick ∼24 non-Rollers that are younger than Rollers and place them onto separate plates. Younger animals among the Roller cohorts can also be picked. For inexperienced injectors, we recommend using 5′ end-modified dsDNA donors and picking F1 Rollers.
4. To avoid false positives due to mosaicism in F1 animals, pick several F2s from each plate, perform pooled lysis and genotype. Genotyping primers should lie outside of the homology arms to avoid amplification from transiently retained donor molecules.
5. Alternatively, correct insertions of fluorescent tags can be screened under a fluorescence dissecting scope or by using high magnification fluorescence microscope. For high magnification screening, mount several F2 animals onto 2% agarose pads and immobilize with levamisole.

## References

Arribere, J. A., R. T. Bell, B. X. Fu, K. L. Artiles, P. S. Hartman et al., 2014 Efficient marker-free recovery of custom genetic modifications with CRISPR/Cas9 in Caenorhabditis elegans. Genetics 198: 837–846.

Brenner, S., 1974 The genetics of Caenorhabditis elegans. Genetics 77: 71–94.

Cho, S. W., J. Lee, D. Carroll, J. S. Kim and J. Lee, 2013 Heritable gene knockout in Caenorhabditis elegans by direct injection of Cas9-sgRNA ribonucleoproteins. Genetics 195: 1177–1180.

Dokshin, G. A., K. S. Ghanta, K. M. Piscopo and C. C. Mello, 2018 Robust Genome Editing with Short Single-Stranded and Long, Partially Single-Stranded DNA Donors in Caenorhabditis elegans. Genetics 210: 781–787.

Ebbing, A., P. Shang, N. Geijsen and H. Korswagen, 2017 Extending the CRISPR toolbox for C. elegans: Cpf1 as an alternative gene editing system for AT-rich sequences, pp.

Farboud, B., A. F. Severson and B. J. Meyer, 2019 Strategies for Efficient Genome Editing Using CRISPR-Cas9. Genetics 211: 431–457.

Forbes, D. J., M. W. Kirschner and J. W. Newport, 1983 Spontaneous formation of nucleus-like structures around bacteriophage DNA microinjected into Xenopus eggs. Cell 34: 13–23.

Ghanta, K. S., G. A. Dokshin, A. Mir, P. M. Krishnamurthy, H. Gneid et al., 2018 5′ Modifications Improve Potency and Efficacy of DNA Donors for Precision Genome Editing. bioRxiv.

Kim, H., T. Ishidate, K. S. Ghanta, M. Seth, D. Conte, Jr. et al., 2014 A co-CRISPR strategy for efficient genome editing in Caenorhabditis elegans. Genetics 197: 1069–1080.

McCarter, J., B. Bartlett, T. Dang and T. Schedl, 1999 On the control of oocyte meiotic maturation and ovulation in Caenorhabditis elegans. Dev Biol 205: 111–128.

Mello, C. C., J. M. Kramer, D. Stinchcomb and V. Ambros, 1991 Efficient gene transfer in C.elegans: extrachromosomal maintenance and integration of transforming sequences. EMBO J 10: 3959–3970.

Paix, A., A. Folkmann, D. Rasoloson and G. Seydoux, 2015 High Efficiency, Homology-Directed Genome Editing in Caenorhabditis elegans Using CRISPR-Cas9 Ribonucleoprotein Complexes. Genetics 201: 47–54.

Paix, A., A. Folkmann and G. Seydoux, 2017 Precision genome editing using CRISPR-Cas9 and linear repair templates in C. elegans. Methods 121-122: 86–93.

Paix, A., H. Schmidt and G. Seydoux, 2016 Cas9-assisted recombineering in C. elegans: genome editing using in vivo assembly of linear DNAs. Nucleic Acids Res 44: e128.

Paix, A., Y. Wang, H. E. Smith, C. Y. Lee, D. Calidas et al., 2014 Scalable and versatile genome editing using linear DNAs with microhomology to Cas9 Sites in Caenorhabditis elegans. Genetics 198: 1347–1356.

Prior, H., A. K. Jawad, L. MacConnachie and A. A. Beg, 2017 Highly Efficient, Rapid and Co-CRISPR-Independent Genome Editing in Caenorhabditis elegans. G3 (Bethesda) 7: 3693–3698.

Schwartz, M. L., and E. M. Jorgensen, 2016 SapTrap, a Toolkit for High-Throughput CRISPR/Cas9 Gene Modification in Caenorhabditis elegans. Genetics 202: 1277–1288.

Silva-Garcia, C. G., A. Lanjuin, C. Heintz, S. Dutta, N. M. Clark et al., 2019 Single-Copy Knock-In Loci for Defined Gene Expression in Caenorhabditis elegans. G3 (Bethesda) 9: 2195–2198.

Stinchcomb, D. T., J. E. Shaw, S. H. Carr and D. Hirsh, 1985 Extrachromosomal DNA transformation of Caenorhabditis elegans. Mol Cell Biol 5: 3484–3496.

Tzur, Y. B., A. E. Friedland, S. Nadarajan, G. M. Church, J. A. Calarco et al., 2013 Heritable custom genomic modifications in Caenorhabditis elegans via a CRISPR-Cas9 system. Genetics 195: 1181–1185.

Vicencio, J., C. Martinez-Fernandez, X. Serrat and J. Ceron, 2019 Efficient Generation of Endogenous Fluorescent Reporters by Nested CRISPR in Caenorhabditis elegans. Genetics 211: 1143–1154.

Ward, J. D., 2015 Rapid and precise engineering of the Caenorhabditis elegans genome with lethal mutation co-conversion and inactivation of NHEJ repair. Genetics 199: 363–377.

Zhao, P., Z. Zhang, H. Ke, Y. Yue and D. Xue, 2014 Oligonucleotide-based targeted gene editing in C. elegans via the CRISPR/Cas9 system. Cell Res 24: 247–250.

